# Graphene-oxide quenching-based molecular beacon imaging of exosome-mediated transfer of neurogenic miR-193a on microfluidic platform

**DOI:** 10.1101/253716

**Authors:** Hyun Jeong Oh, Hyejin Park, Dayoung Yoon, Seok Chung, Do Won Hwang, Dong Soo Lee

## Abstract

A microRNA (miR-193a) was found to be transferred in the exosomes of differentiated neural progenitors to undifferentiated clones. Graphene-oxide (GO) quenching-based molecular beacon was developed to detect RNAs in living cells and tissues quickly and sensitively. Here, we applied GO quencher-based molecular beacon sensor to visualize neurogenic miR-193a levels delivered via exosome during cell-non-autonomous neurogenesis of neural progenitor cells on microfluidic platform. Fluorescence signals of FAM-labeled peptide nucleic acid (PNA) against miR-193a quenched by GO nanosheets (FAM-PNA193a-GO) were recovered in undifferentiated recipient cells differentiated to the neuronal lineage by exosome-mediated neurogenesis 3 days after co-culture with differentiated donor cells. We propose that molecular beacon imaging using PNA-GO complex can be used to visualize individual cellular expression of mature microRNAs revealing their precise spatial localization and temporal sequences by the intercellular exosome delivery of messages to undergo processes such as cell-non-autonomous neurogenesis.

## INTRODUCTION

Neurogenesis is associated with the dynamic changes of microRNA (miRNA) expression and miRNAs are proposed to be responsible for determining cellular status of differentiating neural stem or progenitor cells (*1, 2*). MiRNAs degrade target mRNAs or inhibit translation of mRNAs in a sequence-specific manner (*3-6*) and take roles in cell proliferation (*7, 8*) or differentiation (*9-11*). Several miRNAs such as let-7, miR-124, miR-9 or miR193a are known to be involved in neuronal differentiation of neural stem or progenitor cells (*12-15*).

A neurogenic miRNA was found to regulate neurogenesis by its delivery within exosomes from the donor to the recipient cells (*7, 16*). Exosomes contain many peptides, nucleic acids and miRNAs involved in regulation of proliferation and differentiation (*7, 8, 10, 11, 16-21*). We observed that exosomes from the differentiated neural progenitor cells delivered the neurogenic miR-193a within them to promote neurogenesis of neighboring undifferentiated clones. Functional consequences of miR-193a were verified using bioluminescence imaging using luciferase having sense sequence in their 3’UTR against antisense sequence of miR-193a (*22-24*) using the prototype method to reveal mature miRNA action (*24-26*).

Luciferase reporter imaging is excellent without background even *in vivo* as well as *in vitro*, however, luciferase activity decreases by the miRNA of interest, which is known as a “signal-off system.” Mature miRNA activity is observed by examining the degree of decreased reporter activity. The problem of this signal off system lies in the fact that the reporter signal decrease can also be due to the cell conditions such as dying processes. Molecular beacon was proposed as “signal-on system” for monitoring RNA expression (*27,28*) and we reported the use of fluorescence nanoparticle-based molecular beacon system for monitoring miRNA expression (*29*). Graphene-oxide quenching-based detection system adopted the same signal-on strategy successfully to detect presence of multiplex miRNAs in live cultured cells (*30*).

Neuronal miRNAs turn over rapidly depending in retinal, hippocampal and cortical neurons upon stimuli or activity (*31*) and transported to the dendrites as pre-miRNA to be converted to mature miRNA responding to activity-dependent stimuli (*6, 32, 33*). If a sensing platform is developed for detection of the expression levels of specific mature miRNAs, we would be able to understand the spatially localized and temporarily transient miRNA action in neuronal cell bodies and dendrites.

Graphene oxide and its π-π stacking interaction to absorb and release fluorescent dye-labeled peptide nucleic acid (PNA) probes by hydrogen bonds of endogenous miRNA was used for multiplexed miRNA detection (*30*) and for differential detection of miRNAs showing single-base pair mismatches (*34*). Graphene’s fluorescence quenching capability appears when it attaches fluorescent dye-labeled PNA and fluorescence is recovered according to PNA’s detachment from graphene due to endogenous miRNA having sequences complimentary to PNA probes. PNA, a non-natural nucleic acid analog, has the backbones held by uncharged amide bonds which differs from the negatively charged-phosphodiester bonds of classical nucleic acids (*35, 36*).

The miRNA sensing strategy is based on the fluorescence recovery of the quenched dye-labeled PNA as a probe that was tightly bound to the surface of GO as a fluorescence quencher and subsequent recovery of the fluorescence upon addition of target miRNA (Fig. 1). This miRNA sensing platform allows high sensitivity and specificity toward target miRNA with low background signal. Microfluidic cell culture assay which can be used for observation and capturing of various phenomena between cells, to simulate *in vivo* cellular interaction under *in vitro* setting using tissue-mimetic architectures with hydrogel-incorporating microfluidic device and interstitial mimicking fluid (*37*).

**Fig. 1.**
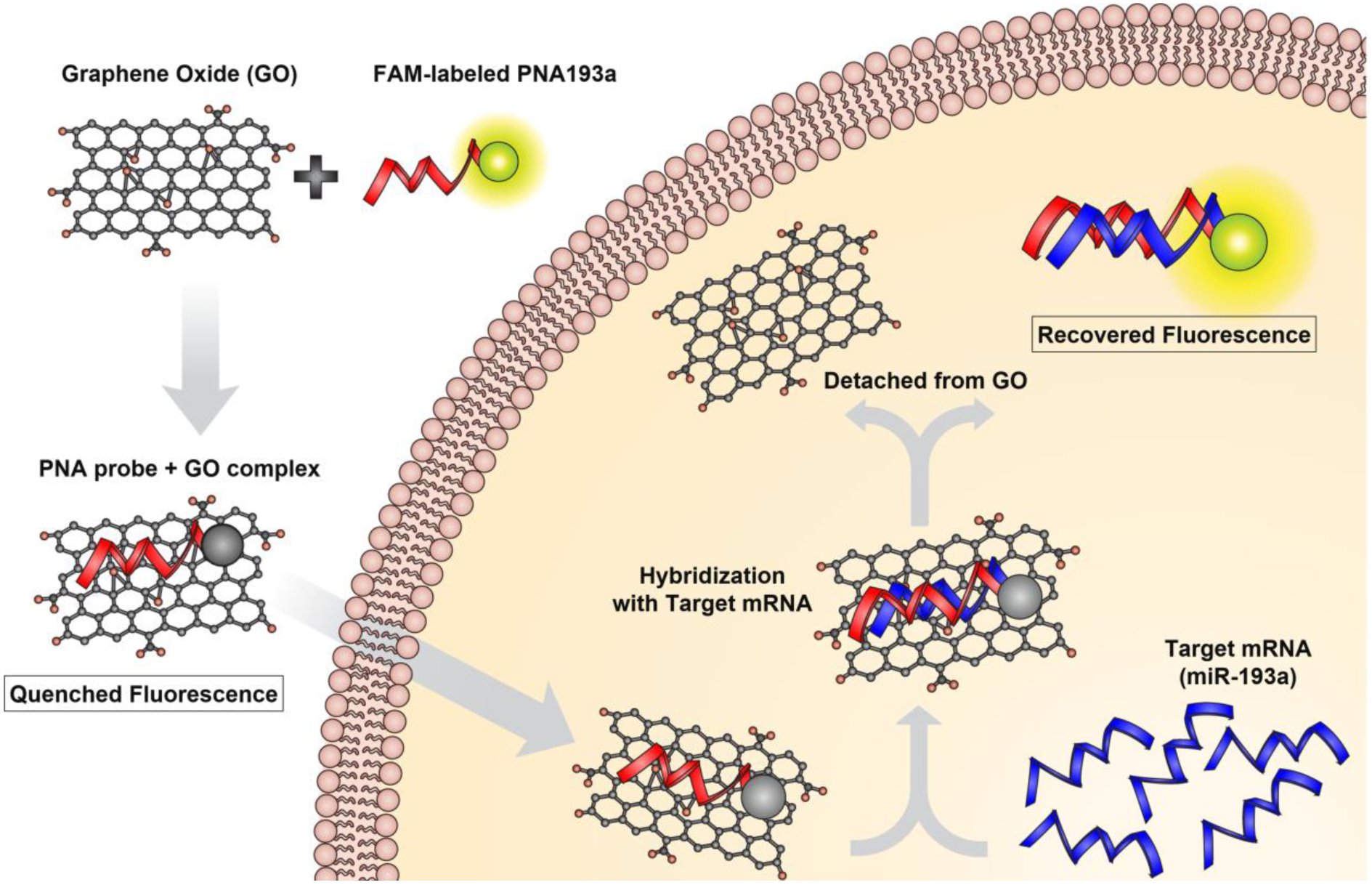
Scheme of strategy for miR-193a sensing based on GO and PNA. The fluorescence signals were recovered when the fluorescent dye-labeled probe was detached from GO by hybridization with a miR-193a in differentiated neural cells.

Here, we applied the above miRNA sensing probes using dye-labeled PNA and GO for sensitive and real-time monitoring of specific miRNAs in living cells on microfluidic devices. We tried to observe individual recipient neural progenitor cells which receives exosomes having relevant miRNAs within and responds to the cell-exosome fusion to start to differentiate to the neurons (*38*). In our previous study (*38*), we established that microfluidic platform could visualize convective exosomal transport and observe exosome-mediated communication between undifferentiated and differentiated cells. Referring to this study (*38*), in this investigation, we established molecular beacon imaging to detect endogenous miRNA expression based on fluorescence dye-labeled PNA-GO beacon during exosome-mediated neurogenesis in the hydrogel-incorporating microfluidic platform.

## Results

### Neurogenesis of neural progenitor F11 cells by miR-193a

In our earlier work, we identified the miR-193a being involved in Ngn1-induced neurogenesis. We validated that miR-193a was highly expressed in differentiated neural cells by Ngn1 or db-cAMP and confirmed the neurogenic function of miR-193a in neural progenitor cells.

To measure miR-193a expression in differentiating neural cells, F11 cells were treated with differentiation media containing db-cAMP for 3 days. Differentiated F11 cells (D-F11 cells) showed a significant neurite outgrowth and the expression of neural marker Tuj-1, but not in undifferentiated F11 cells (UD-F11 cells) (Fig. 2A). On the qRT-PCR, the expression of miR-193a was quantified and high amount of miR-193a was detected only in D-F11 cells (Fig. 2B).

**Fig. 2.**
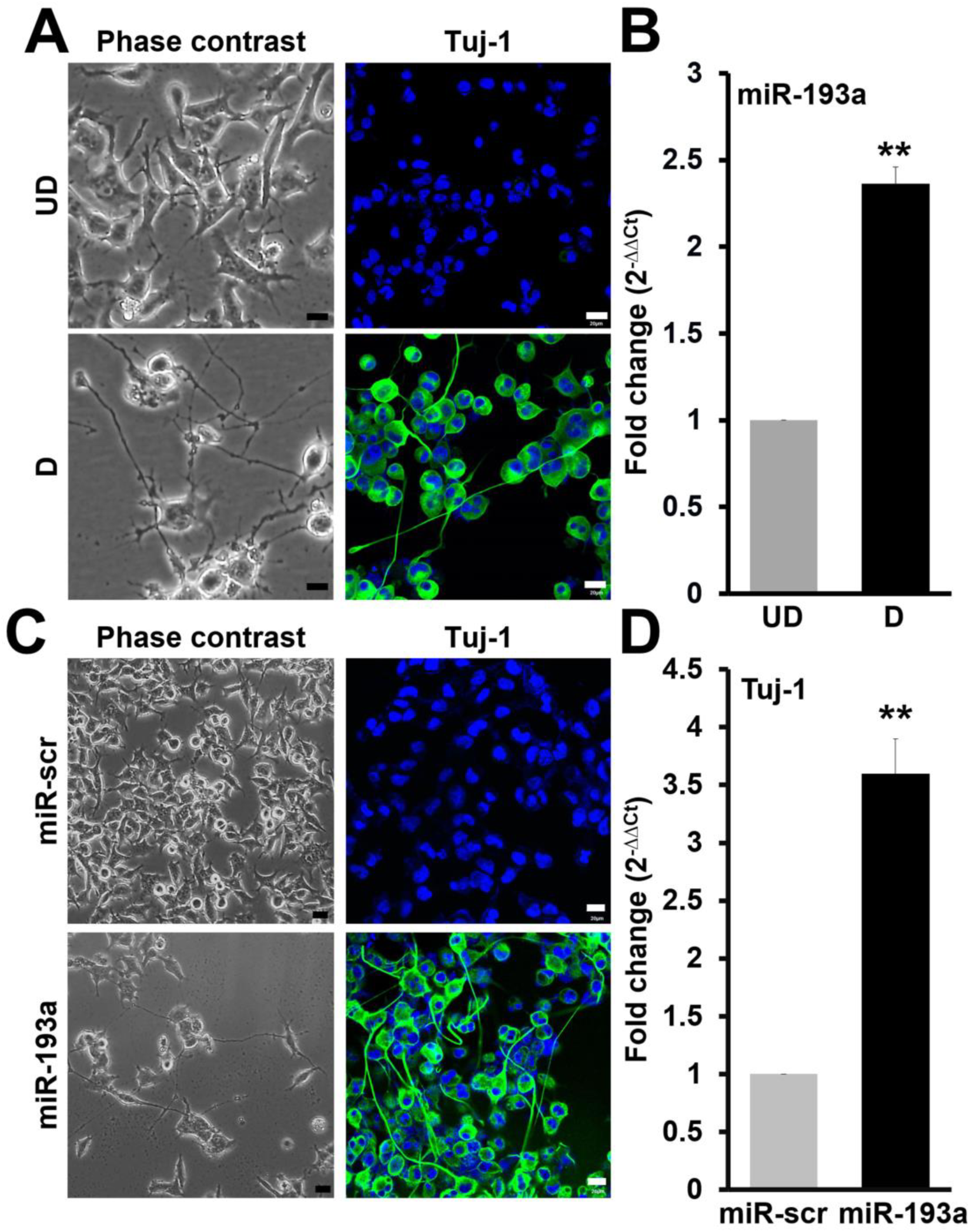
The enhanced expression of miR-193a in cAMP-induced neuronal differentiation of F11 cells. (**A**) Phase contrast images for morphology in UD- and D-F11 cells (Left panel). Scale bar, 10 μm. Representative images of immunofluorescence staining for Tuj-1 in UD- or D-F11 cells (Right panel). Scale bar, 20 μm. (**B**) Quantitative RT-PCR (qRT-PCR) analysis for the expression of miR-193a in UD- and D-F11 cells. Data are representative of three experiments (mean ± s.d). \*\**P* < 0.005. UD: undifferentiated cells, D: differentiated cells. (**C**) Phase contrast images of F11 cells 3 days after transfection with miR-scr or miR-193a (Left panel). Scale bar, 10 μm. Immunofluorescence staining for neural marker, Tuj-1 in F11 cells treated with miR-scr or miR-193a (Right panel). Scale bar, 20 μm. (**D**) qRT-PCR analysis for the expression of neuron-specific marker, Tuj-1 after treatment of miR-scr or miR-193a in F11 cells. **P < 0.005.

To validate the effects of the miR-193a on neurogenesis, F11 cells were transfected with miR-scr or miR-193a for 3 days to induce neurogenesis. The normal F11 cell morphology was maintained and most of the F11 cells underwent proliferation when treated with miR-scr. However, F11 cells transfected with miR-193a showed a significant neurite outgrowth pattern at 3 days after transfection (Fig. 2C) and Tuj-1 expression was examined on immunofluorescence staining (Fig. 2C) and qRT-PCR analysis (Fig. 2D). Tuj-1 expressed highly in F11 cells transfected with miR-193a, compared with that in F11 cells treated with miR-scr (Fig. 2, C and D). In summary, miR-193a was induced during neurogenesis in F11 cells and could induce neurogenesis when transfected.

### FAM-PNA-GO complexes as sequence-specific molecular beacon

We tested the fluorescence quenching of FAM-PNAscr or FAM-PNA193a (fluorescent dye-conjugated PNA probe) by their GO attachment. Fluorescence of FAM-PNAscr or FAM-PNA193a solution (40 pmol) was completely quenched at 0.4 g of GO (Fig. 3A) and we used FAM-PNAscr-GO or FAM-PNA193a-GO mixture at a ratio 40 pmol: 0.4 g for further experiment. PNA193a probe had the complementary sequence to miR-193a. FAM-PNA-GO complex recovered fluorescence by miR-193a but not by miR-scr (Fig. 3B). On cell culture plate, fluorescence signals increased in a miR-193a-concentration dependent manner upon addition of complementary miR-193a (fig. S1).

**Fig. 3.**
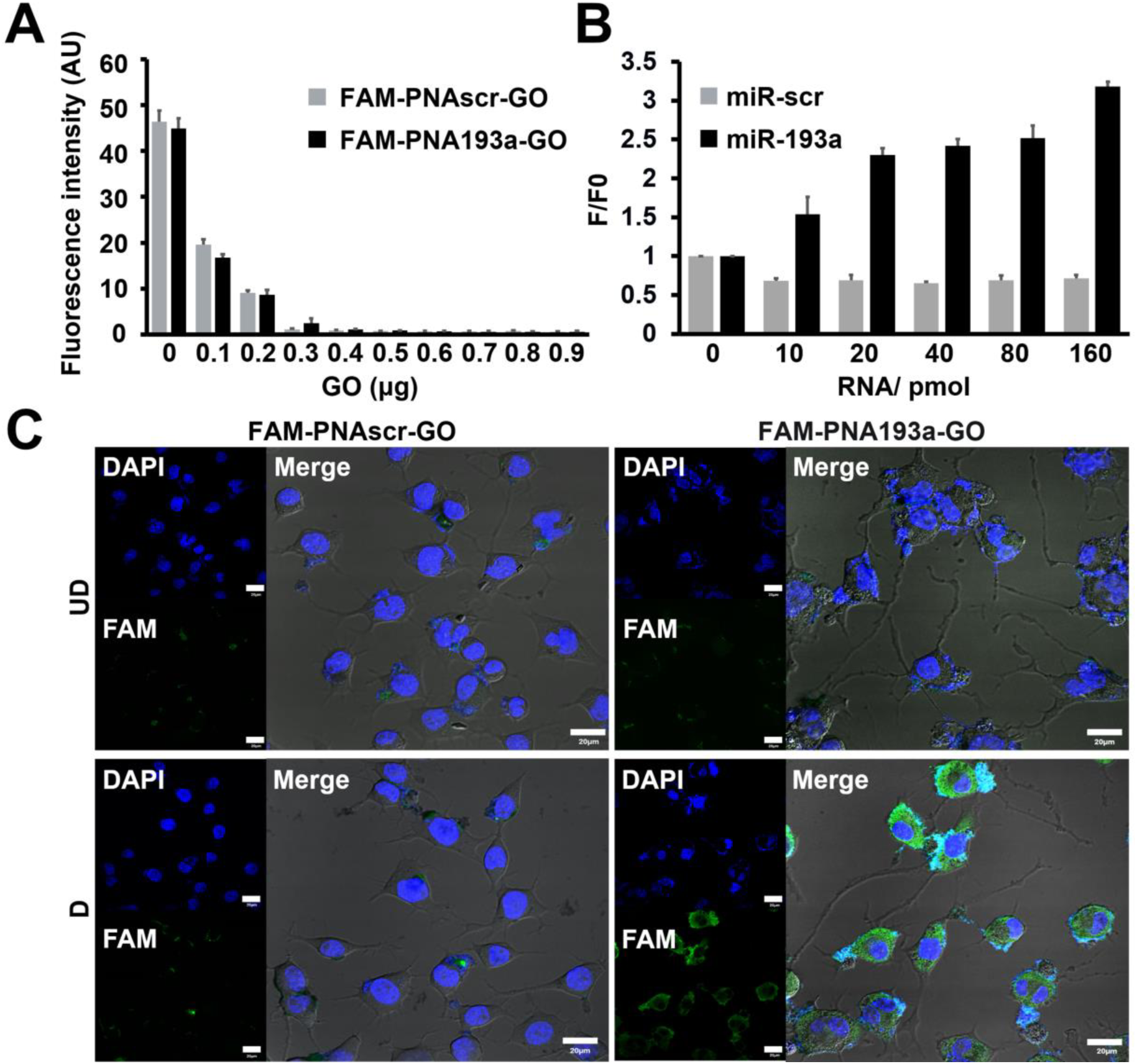
Specificity test of FAM-PNA193a-GO complex against target miR-193a. (**A**) The fluorescence intensity of FAM-PNA193a and FAM-PNAscr (40 pmol) was gradually quenched in a dose-dependent manner after incubation with GO. Fluorescence signals from both FAM-PNA193a and FAM-PNAscr probe were completely quenched with 0.4 μg of GO. (**B**) Quenched FAM-PNA probe-GO complex (40 pmol PNA, 0.4 μg) was mixed with the complementary miR-193a oligomer at different concentration. Fluorescence recovery of the quenched FAM-PNA probe-GO complex was observed upon addition of complementary miR-193a in a concentration-dependent manner, whereas the addition of miR-scr induced no fluorescence recovery. (F, fluorescence intensity of FAM-PNA probe-GO complex in the presence of miR-193a oligomer; F0, basal fluorescence intensity of FAM-PNA probe-GO complex without miR-193a treatment). (**C**) Detection of miR-193a 14 h after treatment of FAM-PNA193a-GO or FAM-PNAscr-GO complex in UD- or D-F11 cells. The intracellularly delivered FAM-PNA193a-GO complex induced the increased fluorescence signals of FAM only in D-F11 cells. Scale bar, 20 μm. UD: undifferentiated cells, D: differentiated cells.

### Visualization of miR-193a expression in differentiating neural progenitor cells using FAM-PNA193a-GO molecular beacon

We examined whether FAM-PNA-GO complex can visualize miR-193a expression in differentiated F11 cells while miR-193a was highly expressed in D-F11 cells on qRT-PCR (Fig. 2B). Addition of GO to FAM-PNA probes quenched fluorescence of FAM completely in opti-MEM medium (fig. S2), and this quenching was not affected in opti-MEM medium while differentiation medium or complete medium increased fluorescence in small amount but significantly by themselves (fig. S3). F11 cells were cultured with differentiation-inducing medium containing db-cAMP for 3 days. The differentiated F11 cells were then incubated with FAM-PNA193a-GO complex for 14 h in opti-MEM medium and after extensive washing fluorescence images of the UD- and D-F11 cells were obtained. As miR-193a was up-regulated during neurogenesis in neural progenitor cells, fluorescence signals were observed in the cytoplasm of D-F11 cells treated with FAM-PNA193a-GO while fluorescence was not recovered in D-F11 cells treated with FAM-PNAscr-GO and in UD-F11 cells treated with FAM-PNAscr-GO or FAM-PNA193a-GO (Fig. 3C). These results indicate that FAM-PNA193a-GO system could clearly distinguish the change of endogenous mature miR-193a expression between pre-and post-neuronal differentiation by sensing the increase in miR-193a level during neurogenesis.

### Molecular beacon imaging of miR-193a expression during exosome-mediated neurogenesis on the microfluidic platform

In our previous study (*38*), we confirmed exosomal transport from differentiated to undifferentiated cells through the exosome-mediated transport of miR-193a in co-culture, transwell culture and on the microfluidic platform. The exosomes were labeled with RFP fused with CD63 to visualize their migration on the microfluidic chambers from undifferentiated to differentiated F11 cells (fig. S4). On this microfluidic system, luciferase-laden recipient undifferentiated neural progenitor cells received exosomes by the convective flow and by showing “signal off” of luminescence showing the functional effect of the miR-193a delivered by exosomes from differentiated cells (*38*).

In the current investigation, we used FAM-PNA193a-GO molecular beacon to monitor the expression of miR-193a by exosome-mediated neurogenesis. We designed a microfluidic platform for the elucidation of functional outcome of exosome-mediated miRNA delivery in recipient chamber with convective flow from the donor chamber containing differentiated neural progenitor cells (Fig. 4A). Donor F11 cells were seeded into the left cell culture channel and cultured with differentiation medium containing db-cAMP to induce neurogenesis. After 48 h, once donor cells were considered to have differentiated, recipient F11 cells were seeded into the right channel. Three days after co-culture in both channels, D-donor cells or recipient cells co-cultured with D-donor cells were treated with FAM-PNA193a-GO molecular beacon for 14 h in opti-MEM medium, and then observed using fluorescence confocal microscopy.

**Fig. 4.**
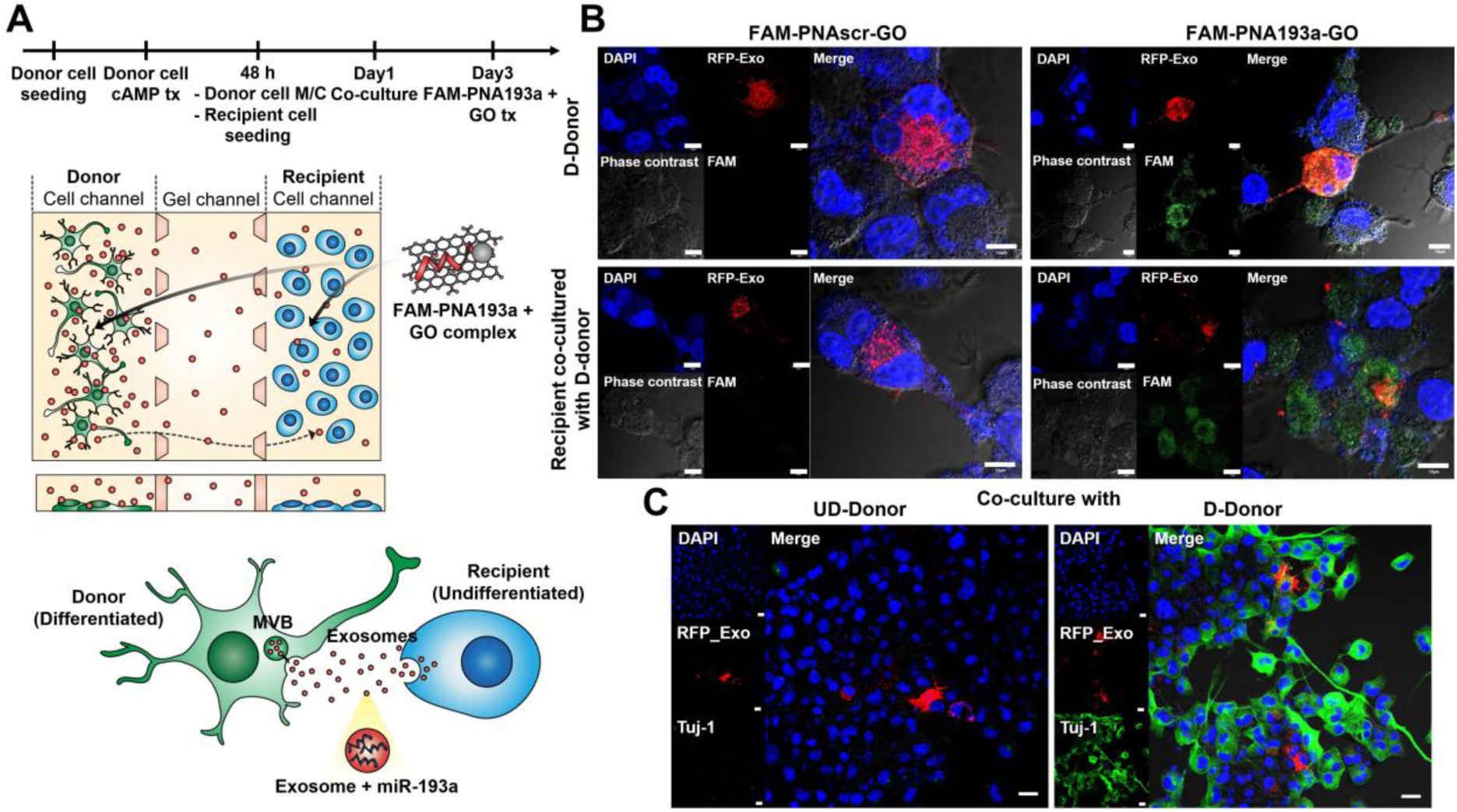
Neurogenic miR-193a sensing during exosome-mediated neurogenesis in the microfluidic assay. (**A**) (Upper panel) Schematic representation of the microfluidic cell culture assay for donor D-F11 cells and recipient UD-F11 cells. Procedure for sequential cell seeding and treatment of FAM-PNA193a-GO complex in the microfluidic assay. (Lower panel) Representative scheme for exosome-mediated miR-193a transfer from donor D-F11 cells to recipient UD-F11 cells. (**B**) Representative fluorescence images showing donor D-F11 cells or recipient F11 cells 14 h after treatment of FAM-PNA193a-GO. Recipient F11 cells were cultured with donor D-F11 cells for 3 days. Fluorescence recovery showing increased expression levels of miR-193a was detected in differentiating recipient F11 cells as well as donor D-F11 cells. Scale bar, 10 μm. (**C**) Immunofluorescence staining for Tuj-1 in recipient F11 cells cultured with RFP-exosome-releasing donor UD- or D-F11 cells. Scale bar, 20 μm.

Fluorescence from D-donor cells and the recipient cells cultured with D-donor cells was recovered from the dark background which represented FAM-PNA193a detachment through the reaction of miR-193a in exosomes of D-donor cells and in the recipient cells having taken up exosomes delivered convectively through the gel channel from the differentiated cells of donor cell channel (Fig. 4B). Fluorescence of recipient cells was only barely seen in recipient cells cultured with undifferentiated donor cells (fig. S5).

### Heterogeneity of exosome-and miR-193a delivery and consequent neural differentiation marker in recipient neural progenitor cells

Exosomes labeled with CD63-RFP released from donor UD- and D-F11 cells moved to and were taken up by recipient F11 cells similarly (fig. S5, red dots). Recipient UD-F11 cells which took up exosomes recovered fluorescence of PNA193a-GO molecular beacon as well as their neighboring cells. On the contrary, some of recipient F11 cells having taken up exosomes did not recover fluorescence. The match (or mismatch) of the cellular exosome uptake and the delivery of miR-193a in those exosomes were well recognized easily by observing the individual recipient F11 cells. In the individual recipient cells, differences in the amount of miR-193a loaded in donor exosomes (or donor exosomes without miR-193a) would have resulted in yielding variable fluorescence recovery. We also found some of recipient cells showing positive fluorescent recovery without taken-up exosomes. Interestingly, recipient F11 cells showed homogeneous Tuj-1 expression in almost all recipient cells regardless of evidence of exosome uptake. In addition to the recipient F11 cells with exosome uptake, other recipient cells without CD63-RFP fluorescence showed homogeneous and even Tuj-1 expression (Fig. 4C). On microfluidic platform with two channels separated by the intervening gel channel, exosome delivery was observed upon convective media flow from donor to recipient channels in either differentiated F11 cells or undifferentiated F11 cells. When the donor cells were differentiated F11 cells, exosome uptake in the recipient cells and consequent miR-193a delivery were heterogeneous but the Tuj-1 expression were homogeneous. Nevertheless, when donor cells were undifferentiated F11 cells neither miR-193a expression nor Tuj-1 expression was observed. Such was also the case when we used PNAscr-GO molecular beacon,

## Discussion

Neurogenic miRNAs play key roles in regulating relevant gene expression for neurogenesis in neural development (*9-11*). Moreover, miRNAs in neurons have a general property of fast turnover and activity plays a regulatory role of miRNA turnover in neuronal cells (*31*). Activity-dependent stimulation of glutamate receptor produced mature miRNA in dendrites as well as in soma in a spatially localized manner (*39*). Therefore, development of highly sensitive sensing platform to detect mature miRNA is needed to understand temporal and spatial expression of mature miRNA if we want to understand the roles of dynamically changing miRNA expression in neuronal cells in physiology and in pathology.

This developmental endeavor ranges from studies of neuronal development or differentiation-related changes to studies of temporal and spatially localized expression of mature miRNAs to take part in various physiologic or pathologic processes. Further to the understanding for the cell-autonomous roles of miRNA in neuronal lineage cells, we also need to elucidate neural cells’ inter-cellular communications either using such small or macromolecules as peptides or lipid mediators or using exosomes containing the package of nucleic acids, peptides or other small molecular mediators. How the inter-cellular communication via exosomes contribute to the differentiation of a cluster of neural stem cells are not understood. To understand this cell-non-autonomous phenomenon, we need the methods of visualizing delivery and action of mediators such as miRNA in individual cells.

In our previous study (*38*), using three methods of 2D co-culture, transwell co-culture and microfluidic assay to investigate donor-recipient communications within clones of neural progenitor cells of different stages of differentiation via exosomes, we proved exosome transfer of miR-193a as intercellular communication mechanism of heralding messages of neuronal differentiation from preceding differentiated cells to the following cells. On microfluidic platform, as we divided firstly differentiated cells and undifferentiated cells in separate channels and made a convective flow of media from donor to recipient cell channels, we confirmed that miR-193a was transferred via the exosomes from differentiated donor cells to the undifferentiated recipient cells to induce neurite outgrowth and the expression of neural markers in recipient cells. The functional outcome of successful miRNA delivery was verified using recipient cells containing transgene of miR-193a-complementary target sequences in its 3’UTR of luciferase transgene. The luciferase activity clearly decreased in recipient undifferentiated cells after receiving exosome-mediated miR-193a on this microfluidic platform (*38*).

However, in this previous study, we needed that neural progenitors should be selected clonally having transgenes integrated into their chromosomes and that the “signal off” nature of luciferase reporter necessitated concomitant verification that the cells were not dying or already dead at most in part. To overcome this inconvenience, the molecular beacon imaging system based on “signal on” methods has been developed. Molecular beacon imaging system was repeatedly reported in the literature (*27-29*), and imaging has the advantage of visualizing individual cell clusters revealing whether their mature miRNAs work in spatial or temporal sequences *in vivo* (*29*) as well as *in vitro* (*30*).

In the current study, we used miRNA sensing method using PNA-GO probe which was used successfully to detect multiple miRNAs in living cells with high sequence specificity and low background signal (*30*). GO’s capacity as delivery vehicle to the cytoplasm, great quenching of attached FAM, and easy detachability of single-stranded nucleic acids attached to them by their complementary sequenced miRNAs allowed multiplexing (*30*) and finding differentially single-base mismatches of target miRNAs (*34*). For these purposes, PNA was reported to work better than classical nucleic acids as their delivery with GO vehicle was easier through cell membrane and their detachability from GO was guaranteed when molecular beacon meets appropriate miRNAs in the cytoplasm.

In this investigation, we used fluorescent dye FAM-labeled molecular beacon, FAM-PNAGO as the imaging tool to visualize mature miRNA action *in vitro*. The PNA-GO probes were reported to minimize nonspecific fluorescence and also provided sensitive and single-base specific response to target mature miRNAs with low background fluorescence signals even in living cells (*30*) and we exploited this advantage for molecular beacon imaging in recipient cells on microfluidic platform.

The microfluidic cell culture platform enabled us to observe cell-cell interaction at high resolution in real time while allowing easy modification or controlling the sophisticated fluidic system (*37*). We used the same miRNA-sensing microfluidic platform as our previous study (*38*) but with “signal-on” molecular beacon of FAM-labeled PNA-GO complex for the detection of the presence of neurogenic mature miR-193a delivered by exosomes while varying interstitial flow to simulate *in vivo* situation. This microfluidic platform allowed sequential seeding and precedent induction of neurogenesis for donor cells and observation of recipient cells to receive exosomes and their contents. On this microfluidic channels, both the donor and the recipient cells were viable by refreshing medium daily, compared to conventional transwell co-culture studies (*37, 38*). Furthermore, this microfluidic platform allowed repeated observation of the recipient channel cells using usual confocal microscopes and thus enabled the visualization of molecular beacon “signaling on” within the recipient cells.

As miR-193a was expressed during neurogenesis and was transferred via exosomes from differentiated cells to undifferentiated cells on microfluidic platform, we used FAM-PNA193a-GO as molecular beacon for imaging on microfluidic platform. On this platform, miR-193a was highly expressed in recipient cells at all 3 days of co-culture, but neurite outgrowth started 2 days after co-culture and neural marker appeared 3 days after co-culture (*38*). These results indicate that exosomal miR-193a from differentiated donor cells induces neurogenesis of undifferentiated recipient cells at first and this promotes action of miR-193a in recipient cells which took up exosomes, which triggers sequential process of neurogenesis. However, the further studies need to be performed to find out whether the recipient cells having taken up exosomes differentiate alone or the adjacent cells differentiate to, or whether neurogenic miR193a-induced differentiating neurons begin to differentiate to neurons and recover fluorescence as secondary (or tertiary) endogenous miR-193a expression. In the Fig. 4B, CD63-RFP-bound exosomes derived from donor cells were taken up by a certain portion of recipient cells, which showed recovery of FAM fluorescence meaning the presence of mature miR-193a. Interestingly, the cells adjacent to the differentiation-heralding cells which received CD63-RFP exosomes also showed recovery of FAM fluorescence. We speculate the exosomal delivery of neurogenic miR-193a starts differentiation in the recipient cells but also delivers the same or other neurogenic messages to the adjacent cells in a cell-non-autonomous way. Again, unexpectedly, the neuronal marker expression of the recipient cells in Fig. 4C was more extensive in the adjacent cells even having not received any exosomes from donor cells (absence of CD63-RFP). This implies there might be cascade delivery between undifferentiated cells in the recipient channel. Though we did not prove whether the secondary transfer from CD63-RFP positive to CD63-RFP negative recipient cells are also mediated by exosomes, our results would have been observed if the cascade of neurogenesis occurs via miR-193a containing exosomes.

We propose that FAM-PNA-GO probe can act as molecular beacon imaging tool capable of detecting the presence of miR-193a during neurogenesis in neural progenitor cells in cell-non-autonomous fashion. Fluorescence of quenched FAM-PNA193a-GO probe was not recovered by miR-scr oligomer (Fig. 3B). Fluorescence recovery of FAM-PNA193a-GO with matching complimentary miRNAs was dose dependent. Transmembrane cytoplasmic uptake of molecular beacon made of GO and PNA seemed very homogenous as we observed fluorescence recovery in every cell of cAMP-stimulated differentiated clones (Fig. 3C, lower right picture). Obviously, background was very dark when the donor cells were undifferentiated cells though we could observe CD63-RFP in the recipient cells. When the donor cells were differentiated cells but the molecular beacon was FAM-PNAscr-GO with scramble sequences, we observed no fluorescence recovery in the recipient cells regardless of CD63-RFP signals in the cells indicating the presence or absence of exosome delivery (fig. S5). We propose that this molecular beacon of FAM-PNA193a-GO can be used for visualization of individual cells’ response to the exosome-mediated miRNA delivery and the recipient cells’ response to this transfer in a sensitive and specific way with nearly absent background in living cells.

Heterogeneity of exosome access to the cells despite the assumed ready uptake by the recipient cells, and temporally sequential transfer of differentiation message to the adjacent cells from this first encountering cells imply the possibility of understanding what might happen *in vivo* situation. Precedent differentiating cells responding to internal clock or certain external stimulus will recruit colleague cells using exosomal delivery of appropriate message to the adjacent cells to follow the differentiation process in sequential manner. This will make a gradient of apparently synchronous differentiation process and the individual cellular responses to the messages received from the “donor” cells might look heterogeneous. The blotting or qRT-PCR or whichever assay using tens or thousands of cells cultured *in vitro* will limit to elucidate this kind of information regarding how cells change information of what to do next.

Individual miRNAs in neurons turn over rapidly in physiologic or pathologic conditions in dendrites of cerebral neurons (*39*) or in axons of peripheral nerves (*31*). This fast turnover was observed in neuron-enriched miRNAs such as miR-9, miR-125b, and miR-146a have half-lives of 1 to 3.5 h (*40*). The rapid turnover of neuron-specific miRNAs was dependent on neuronal activity (*31*). And neurons in hippocampus activity-dependent maturation of pre-miRNA in the dendrites responding to the stimulus activity (*39*). Based on these findings, localized miRNA maturation was proposed to modulate target gene expression with spatial and temporal precision. We propose our molecular beacon imaging using fluorescent dye-PNA (against a specific miRNA)-GO complex as probes while the cellular uptake is perfect but the sequential imaging of the cells of interest shall reveal the temporal and spatial sequences of intercellular message transfer during physiologic or pathologic processes.

The important caveats of using our GO quenching-based molecular beacon imaging for characterizing the responses of individual cells at single cell level, are 1) sufficient amount of GO molecular beacon should be administered so that the recovered fluorescence be homogeneous, 2) cells shall be located in the recipient chamber of the microfluidic device and should be able to be discerned one by one, and 3) more desirably, dose response relationship according to the amount of a specific active mature microRNA allow characterizing the quantity (activity) of the microRNAs contained in the exosomes.

Secondary or tertiary or further signal transfer between recipient cells stimulates our curiosity as differentiation marker indicates the entire cells eventually participated in the differentiation, and thus further devising a refined platform will be needed to dissect this cell-cell interaction. Homogeneous Tuj-1 expression of secondary recipient cells provokes speculation regarding the plausible spectrum of phenomena that exosomes and their contents released from recipient differentiation-beginning cells mediate further the secondary outcomes or that other totally new biomolecule mediators would have mediated this signal transfer to result in secondary Tuj-1 expression. Though we used the clones of preceding differentiating cells and undifferentiated cells of the same clones, the microfluidic platform will allow different tissue cells of different germinal origins.

## Materials and Methods

### Cell culture

F11 cells, rat dorsal root ganglion and mouse neuroblastoma hybrid cells were cultured in Dulbecco’s modified Eagle’s medium (DMEM, Gibco) supplemented with 10% fetal bovine serum (FBS; Gibco), 10 U/ml penicillin, and 10 μg/ml streptomycin in a humidified atmosphere of 5% CO_2_ at 37 °C. To induce neuronal differentiation, F11 cells were incubated with DMEM containing 0.5% FBS and 1 mM dibutyryl cyclic AMP (db-cAMP, Sigma-Aldrich) or transfected (Lipofectamine2000; Invitrogen) with miR-193a (Ambion^®^) for 3 days.

### Synthesis of PNA and DNA oligomer

The fluorescent dye (FAM)-labeled PNA probe was synthesized by Panagene Inc. The sequences of oligomers were 5′-ACTGGGACTTTGTAGGCCAGTT-3′ (FAM-PNA193a complementary oligomer) and 5′- TTGACCGGATGTTTCAGGGTCA-3′ ′ (FAM-PNAscr complementary oligomer). All PNA probes consisted of 22-mer in length (table 1).

### Quantitative RT-PCR analysis

Total RNA from differentiated F11 cells was prepared using Trizol (Invitrogen) and mirVana^TM^ miRNA Isolation Kit (Ambion^®^). Isolated RNA was analyzed for purity and concentration in a Nanodrop-1000 Spectrophotometer (Thermo Scientific).

The cDNA samples for miRNA was prepared by reverse transcribed (RT) with the miRNA 1st-strand cDNA synthesis kit (Agilent Technologies) and RT-PCR amplification was performed using ABI^®^ 7500 (Applied Biosystems^TM^) with the TaKaRa SYBR Green Master mix (Clontech Laboratories), which is specific for mature miRNA sequences.

PCR primer for miR-193a was 5′-AACTGGCCTACAAAGTCCCAGT-3′ (forward) and the universal reverse primer was used for reverse primer. Relative value normalized to U6 snRNA as an internal control. All experiments were performed in triplicate.

### MiR-193a detection using GO-PNA complexes

FAM-PNA probe (40 pmol per probe) was mixed with a series of GO (Lemonex Inc) solution, concentration ranging from 0.1 μg to 0.9 μg in 100 μl of buffer (Tris-HCl, pH 7.5) for 15 min at room temperature. The quenched fluorescence signals were monitored after the formation of FAM-PNA probe-GO complex. FAM-PNA-GO complex solution were mixed with various concentration (2.5-40 pmol in 50 μl of buffer (Tris-HCl, pH 7.5) of target oligomer having a complementary sequence. Fluorescence signals were measured using a Varioskan Flash Multimode Reader (Thermo Fisher Scientific). The fluorescence images of mixtures in a 96-well black plate were obtained using IVIS-100 imaging system (Xenogen).

### MiR-193a detection in differentiated cells

F11 cells were seeded on 6-well plate at 1.5 × 10^5^ cells per well and incubated with DMEM containing 0.5% FBS and 1 mM db-cAMP (Sigma-Aldrich) for 3 days. FAM-PNA probe (100 pmole per probe) was mixed with the GO (1 g) in opti-MEM media (Invitrogen) for 15 min at room temperature. The FAM-PNA193a-GO complexes were treated to F11 cells for 14 h at 37 °C and then vigorously washed three times with PBS. The cells were fixed with 4% paraformaldehyde (PFA) and nuclei were counterstained with 4’-6-diamidino-2-phenylindole (DAPI, Vector Laboratories). Fluorescent images were obtained using a FV1000 confocal laser scanning microscope (Olympus).

### Fluorescent labeling of exosomes

F11 cells (1.5 x 10^5^ cells per well) were plated in 6-well plates and transfected with CMV-driven copepod GFP-tagged CD63 vector (System Biosciences) using Lipofectamine2000 (Invitrogen) diluted in OPTI-MEM medium (Gibco). The transfected F11 cells were incubated in DMEM supplemented with 10% FBS or 0.5% FBS and 1 mM db-cAMP for 2 days. Fluorescent images were analyzed using a FV1000 confocal laser scanning microscope (Olympus).

### Microfluidic cell culture assay

#### Microfluidic Device Fabrication

The microfluidic co-culture system is composed of polydimethylsiloxane (PDMS; Sylgard 184; Dow Corning). The SU-8 photoresist pattern master (MicroChem) was used as a master mold, using a conventional soft lithography process to replicate the microchannel-patterned PDMS. PDMS mixed with the curing agent at a 10:1 weight ratio was poured onto the wafer and cured by baking in an oven at 60 °C for 8 h. PDMS replica was detached from the wafer and all reservoir patterns of the PDMS replica were punched by dermal biopsy punches (a 6 mm punch for media reservoirs and a 1 mm punch for gel filling reservoirs). The PDMS replica and glass coverslip were sterilized and bonded together via oxygen plasma (Femto Science) and placed at 60 °C in an oven for at least 48 h to restore hydrophobicity of the microchannel surfaces.

#### Co-culture of F11 cells in the microfluidic system

Type 1 collagen ECM (2 mg/ml; BD Biosciences) was diluted to 2 mg/mL in a mixture of 10 × phosphate-buffered saline (PBS; Gibco) and distilled deionized water. The pH of the hydrogel solution was optimized to 7.4 with 0.5 N NaOH. Type 1 collagen ECM was inoculated into hydrogel channels and gelled for 30 minutes. The cell culture channel was then filled with medium to prepare cell seeding. F11 cells (donor cells) were seeded into the cell culture channel (left channel) and conditioned medium was added into the right cell culture channel. After cell attachment, left channel containing donor cells of the differentiated group, the medium was replaced with differentiation medium (DMEM containing 0.5% FBS and 1 mM db-cAMP) for 48 h. F11 cells (recipient cells) were seeded into cell culture channel (right) and co-culture with donor cells at 37 °C in 5% CO_2_ atmosphere for 3 days. Density of F11 cells (donor and recipient) suspended in conditioned medium was 1 × 10^6^ cells/mL.

Three days after co-culture, FAM-PNA probes (100 pmole per probe) mixed with the GO (1 g) in opti-MEM media (Invitrogen) were incubated for 15 min at room temperature. The FAM-PNA193a-GO complexes were treated to recipient F11 cells (right cell culture channel) for 14h at 37 °C and then vigorously washed three times with PBS. The cells were fixed with 4% paraformaldehyde (PFA) and nuclei were counterstained with 4’-6-diamidino-2-phenylindole (DAPI, Vector Laboratories). Fluorescent images were obtained using a FV1000 confocal laser scanning microscope (Olympus).

### Statistical analysis

Results of all experiments were collected from three or four independent experiments for each sample. Data are presented as means ± standard deviation (SD). The Student’s *t*-test was used to calculate P values. Statistical significance was accepted at *P-*values of < 0.005.

## Supplementary Materials

fig. S1. Specificity test of FAM-PNA probe-GO complex after treatment of complementary MIR-193a oligomer.

fig. S2. Quenched fluorescence intensity of FAM-PNA probes in opti-MEM.

fig. S3. Recovered fluorescence intensity of FAM-PNA probe-GO complex after incubation with medium.

fig. S4. Fluorescent labeling of exosomes.

fig. S5. Neurogenic miR-193a sensing during exosome-mediated neurogenesis in recipient cells cultured with UD- or D-donor F11 cells.

table S1. Sequence information of peptide nucleic acid (PNA) probe for miR-193a. The miR-scr was used as a negative control.

## Acknowledgements

This research was supported by Basic Science Research Program through the National Research Foundation of Korea(NRF) funded by the Ministry of Education (NRF-2017R1A6A3A01007423), and Radiation Technology R&D program through the National Research Foundation of Korea founded by the Ministry of Science, ICT & Future Planning (NRF-2017M2A2A7A02019899), and by the National Research Foundation of Korea (NRF-2017R1A2B3007701) grant funded by the Korea government(MEST), and by a grant of the Korea Health Technology R&D Project through the Korea Health Industry Development Institute (KHIDI), funded by the Ministry of Health & Welfare, Republic of Korea (HI14C3344, HI14C1277), and National Research Foundation of Korea grant funded by the Korea government (MSIP) (2015M3C7A1028926, 2017M3C7A1048079). **Author Contributions:** H.J. Oh, D.W. Hwang and D.S. Lee conceived and designed the experiments. H.J. Oh performed the experiments. H.J. Oh, D.W. Hwang, D.S. Lee, and S. Chung analyzed the data. H.J. Park, D.Y. Yoon and S. Chung contributed reagents/materials/analysis tools. H.J. Oh, D.W. Hwang and D.S. Lee wrote the manuscript. **Competing interests:** The authors declare that they have no competing interests. **Data and materials availability:** All data needed to evaluate the conclusions in the paper are present in the paper and/or the Supplementary Materials. Additional data related to this paper may be requested from the authors.

